# ATF5-Dependent GDF15 Expression Mediates Anesthesia-Induced Neuroprotection Against Stroke

**DOI:** 10.1101/2024.12.06.627155

**Authors:** Xianshu Ju, Tao Zhang, Jianchen Cui, Yulim Lee, Suho Lee, Ho Min Kim, Boohwi Hong, Jiho Park, Chul Hee Choi, Hyon-Seung Yi, Jun Young Heo, Woosuk Chung

## Abstract

The incidence of perioperative stroke, a rare but severe complication, is increasing in aging populations. Although anesthetics such as sevoflurane may provide protective preconditioning against ischemic injury, clinical outcomes have been inconsistent. In this study, we demonstrated that sevoflurane-induced neuroprotection is associated with the upregulation of genes involved in the mitochondrial unfolded protein response (UPR^mt^) and mitochondrial bioenergetic metabolism. Our findings emphasize the critical role of ATF5, a key transcription factor, in mediating these protective effects. Specifically, we observed that sevoflurane preconditioning significantly upregulates ATF5 and its downstream target GDF15 – a regulator of mitochondrial function – in the cerebral cortex. Notably, we found that this mechanistic pathway was not activated in the brain of aged mice, suggesting that age-specific strategies may be necessary for reducing the risk of perioperative stroke. Considering the steadily increasing age of patients, therapeutic approaches that enhance mitochondrial function in the aged brain may provide additional protection against perioperative stroke.

## Introduction

Perioperative stroke, a term applied to strokes that occur during or within 30 days of surgery, is a relatively rare but devastating perioperative complication (Fanning *et al*, 2024). Although uncommon (0.03-4%) after surgeries other than cardiac surgery and neurosurgery (Ng *et al*, 2011), the incidence of perioperative stroke is increasing in aging societies (Al-Hader *et al*, 2019; Marcucci *et al*, 2023; Smilowitz *et al*, 2017). Studies have reported significantly worse prognosis after perioperative ischemic strokes compared with community-onset strokes, as reflected in increased 30-day mortality, longer hospital stays, increased long-term cognitive impairment and disability (Fanning *et al*., 2024). Recently, the significance of covert perioperative strokes (silent brain infarcts, strokes detectable only by imaging) have also been recognized (Fanning *et al*., 2024). Not only is their incidence much higher (7%), covert strokes are also associated with increased risk of postoperative delirium, cognitive decline, and additional cerebrovascular insults 1 year after surgery (Neuro, 2019). Considering that 310 million individuals worldwide are estimated to undergo surgical procedures annually (Rupert M. Pearse, 2016), there is an imperative need to identify possible treatment measures that reduce the negative impacts of perioperative stroke.

One potential strategy is anesthetic preconditioning (APC), an approach based on the observation that exposure to volatile anesthetic agents causes a transient increase in tolerance against ischemia/reperfusion injury (Lan Wang, 2008; Swyers *et al*, 2014). Preclinical studies have demonstrated that exposure to volatile anesthetics before an ischemic event provides neuroprotection (Hoffmann, 2016; L. Tong, 2014; Qianzi Yang, 2012; Ye *et al*, 2012). They have also shown that sevoflurane, currently the most widely used volatile anesthetic agent, can reduce neurological severity, cerebral infarct size, and brain water (Deng *et al*, 2020). Despite promising findings in preclinical studies, clinical trials have yielded inconsistent results (Lomivorotov *et al*, 2022; Raub *et al*, 2021). However, the consistent neuroprotective effects in preclinical studies are sufficiently compelling to suggest that understanding the mechanism underlying sevoflurane-induced preconditioning in animals might provide important insights that could aid in parsing the results of clinical studies and discovering potential targets for mitigating perioperative stroke.

Mitochondria dysfunction, a hallmark of ischemia/reperfusion injury, leads to neuronal death, highlighting the importance of preserving mitochondrial function for neurological recovery (Peter S. Vosler, 2009; Yang, 2018). Interestingly, mitochondria are also integrally involved in anesthesia-induced preconditioning (Halestrap *et al*, 2007; McLeod *et al*, 2004; Venkatesh *et al*, 2019; Wang *et al*, 2019; Xiao *et al*, 2011; Ye *et al*., 2012). Although the mechanisms underlying mitochondrial protection are not fully understood, recent studies suggest involvement of the mitochondrial unfolded protein response (UPR^mt^) (An *et al*, 2024; Venkatesh *et al*., 2019; Wang *et al*., 2019). UPR^mt^, a component of the integrated stress response, is a transcriptional response that serves to restore mitochondrial function and cellular homeostasis (Anderson & Haynes, 2020; Melber & Haynes, 2018; Zhang *et al*, 2024). A previous study showed that ischemia-induced preconditioning failed to develop in cardiac tissue of transgenic mice lacking the transcriptional factor ATF5 (activating transcription factor 5), a major regulator of UPR^mt^ in mammalian cells (Wang *et al*., 2019). A more recent study demonstrated that pharmacological treatment with meclizine, which modulates mitochondrial respiration and reduces oxidative stress through ATF5-dependent UPR^mt^, provides neuroprotection against ischemia/reperfusion injury (An *et al*., 2024). Since sevoflurane exposure also increases the expression of ATF5 (Lee *et al*, 2021), it is highly possible that ATF5-dependent UPR^mt^ is also intimately involved in sevoflurane-induced preconditioning in the brain.

To confirm the significance of ATF5-dependent UPR^mt^ in sevoflurane-induced preconditioning, we first exposed transgenic mice lacking ATF5 specifically in cortical excitatory neurons (Emx1-Cre;ATF5^fl/fl^ mice) to sevoflurane and then performed a middle cerebral artery occlusion (MCAO) procedure. After confirming the absence of sevoflurane-induced preconditioning, we next investigated the expression of genes related to mitochondria energy metabolism and UPR^mt^ after sevoflurane exposure to identify ATF5-downstream mediators. Interestingly, we found a significant increase in GDF15 (growth differentiation factor 15), a representative mitokine known to regulate energy metabolism in response to ischemia/reperfusion injury in the heart (Hyo Kyun Chung, 2017; Schindowski *et al*, 2011; Yatsuga, 2015; Zhang *et al*., 2024; Zhang *et al*, 2017). By overexpressing ATF5 specifically in excitatory neurons and studying GDF15 whole-body knockout (KO) mice, we provide direct evidence that sevoflurane exposure induces ATF5-dependent release of GDF15 in excitatory neurons, resulting in increased mitochondrial function and resistance against ischemic/reperfusion injury. Importantly, we also found that sevoflurane-induced activation of the ATF5- GDF15 signaling pathway is absent in aged mice, providing a plausible explanation for the inconsistencies between preclinical and clinical studies regarding anesthesia-induced preconditioning.

## Results

### Sevoflurane-induced neuroprotection involves upregulation of genes associated with mitochondrial metabolism and UPR^mt^

To elicit sevoflurane-induced neuroprotection, we exposed mice to 2.5% sevoflurane for 2 hours in an anesthesia chamber. Blood gas levels measured at the end of anesthesia showed that sevoflurane did not cause respiratory depression (Table 1). Twenty-four hours after sevoflurane exposure, cerebral ischemia was induced by MCAO (Fig. 1 A). Proper positioning of the monofilament at the MCA was confirmed using laser Doppler microscopy (Fig. 1 B). Consistent with previous studies, sevoflurane preconditioning reduced infarct size in the cortex and improved neurological scores (Fig. 1 C, D). To evaluate the possible role of UPR^mt^, we next investigated expression levels of representative UPR^mt^ proteins (Fig. 1 E, Suppl. Fig. 1). We found significant increases in the expression of the protease, LONP1, and the transcriptional factor, ATF5. The fact that ATF5 acts as a major regulator of UPR^mt^ (Wang *et al*., 2019) suggests the possibility that ATF5-dependent UPR^mt^ mediates sevoflurane-induced preconditioning by enhancing mitochondrial function in the brain. However, there was no significant increase in mitochondrial respiration in mitochondria isolated from the cerebral cortex (Fig. 1 F), indicating that sevoflurane-induced ATF5 expression may contribute to preconditioning not by increasing mitochondrial activity but by preserving mitochondrial function.

**Figure 1.**
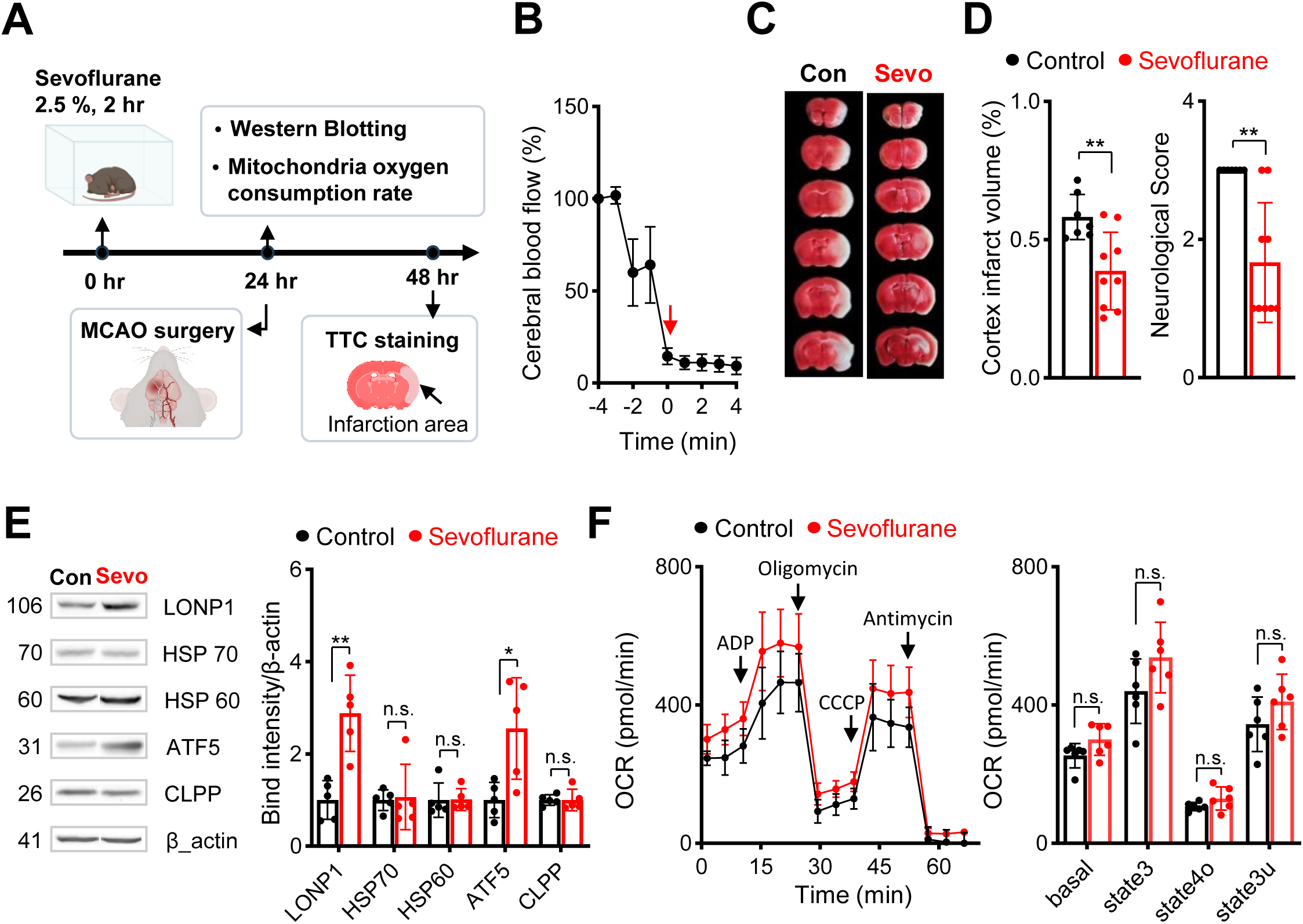
Sevoflurane exposure provides neuroprotection against ischemia/reperfusion injury and upregulates UPR^mt^ proteins. **(A)** Experimental timeline. **(B)** Reduction in cerebral blood flow in the MCA territory during the ischemic period (n = 6). **(C)** Representative TTC-stained images after MCAO in Control and Sevoflurane groups. **(D)** Sevoflurane-induced reduction in cortical infarct volume and improved neurological scores, n = 7-9 per group. **(E)** Western blot analysis of cortex samples obtained 24 hours after sevoflurane exposure, n = 5 per group. **(F)** Left: Mitochondrial function, measured as mitochondrial OCR, determined by assessing mitochondrial respiration in mitochondria samples isolated from the cortex. ADP, oligomycin, CCCP, and antimycin were added sequentially, as indicated by arrows. Right: Quantification of OCR after excluding non-mitochondrial respiration, n = 6 per group. Full Western blot images are provided in Suppl. Fig 1. Values are presented as means ± SD (n.s., not significant; *p < 0.05, **p < 0.01,***p < 0.001).

**Table 1.** Analysis of blood gases and lactate after exposure to 2.5% sevoflurane for 2 hours.

|  | pH | pCO<sub>2</sub><br>(mmHg) | pO<sub>2</sub> (mmHg) | sO<sub>2</sub> (%) | Lactate<br>(mmol/L) |
| --- | --- | --- | --- | --- | --- |
| Animal 1 | 7.375 | 34.3 | 109 | 98 | 1.11 |
| Animal 2 | 7.332 | 38.4 | 85 | 96 | 0.99 |
| Animal 3 | 7.346 | 41.7 | 91 | 97 | 1.19 |
| Mean ± SD | 7.35 ± 0.18 | 38.1 ± 3.0 | 95 ± 10 | 97 ± 0.8 | 1.10 ± 0.08 |

pCO<sub>2</sub>, partial pressure of carbon dioxide; pO<sub>2</sub>, partial pressure of oxygen; sO<sub>2</sub>, oxygen saturation

### Sevoflurane-induced preconditioning does not develop in mice with reduced ATF5 expression in excitatory neurons

To evaluate the significance of ATF5 in excitatory neurons during sevoflurane preconditioning, we generated *Atf5*-cKO (Emx1^cre/+^;*Atf5*^f/f^) mice by crossing Emx1^cre/+^ knock-in mice with *Atf5*^f/f^ mice (Suppl. Fig. 2). RT-qPCR analyses confirmed a significant reduction in the transcription levels of total *Atf5* and its two splice variants (*Atf5*-α and -β) in samples obtained from the cerebral cortex (Fig. 2 A). Because Cre-mediated recombination in this model is limited to excitatory neurons, we further evaluated ATF5 expression in cultures of primary cortical neurons obtained from *Atf5*-cKO mice (Fig. 2 B). Unexpectedly, we found that *ATF5* expression was decrease by only 30% in excitatory neurons, identified by their expression of CaMKII (calcium/calmodulin-dependent protein kinase II) (Fig. 2 B). Immunohistochemistry also confirmed residual ATF5 expression in excitatory neurons in the cerebral cortex of *Atf5*-cKO mice (Fig. 2 C). Notably, sevoflurane failed to reduce infarction size or improve neurological scores in MCAO model *Atf5*-cKO mice, despite the incomplete knockout of *Atf5* (Fig. 2 D, E). These results confirm the significant role of ATF5 in sevoflurane-induced preconditioning in the brain.

**Figure 2.**
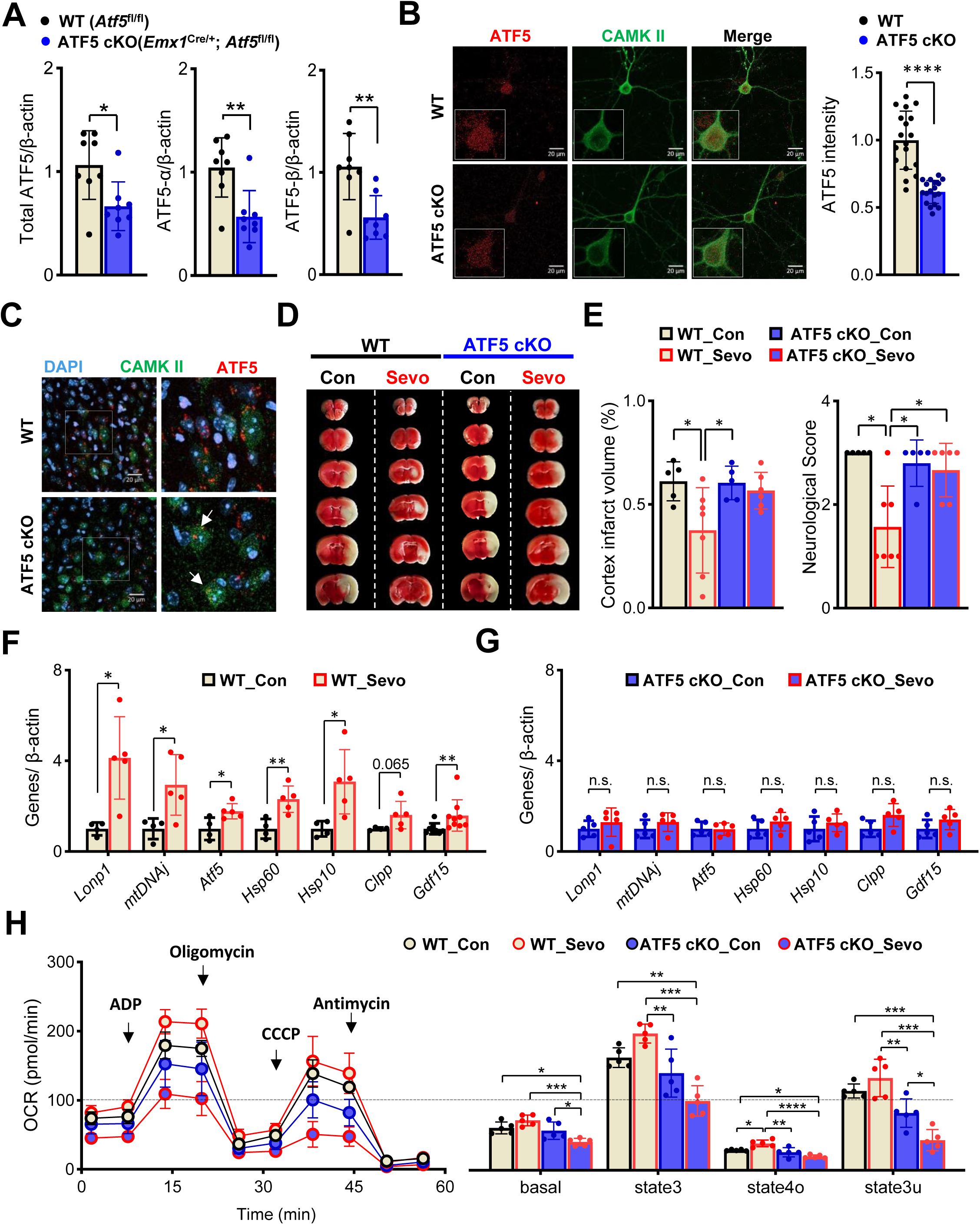
Sevoflurane exposure does not induce preconditioning in *Atf5*-cKO mice but does induce mitochondrial dysfunction. (A) mRNA expression levels of total *Atf5*, *Atf5*-α, and *Atf5*-β in the cortices of WT and *Atf5*-cKO mice, n = 8 per group. (B) Representative immunocytochemical images of cultured primary neurons, n = 18 per group. (C) Representative IHC images of anterior cingulate cortex layer V-VI of *Atf5*-cKO mice showing the colocalization of DAPI (blue), CAMKII (green) and ATF5 (red). White arrow indicates decreased, but remnant, ATF5 expression in excitatory neurons. (D) Representative TTC-stained images of MCAO model WT and *Atf5*-cKO mice after sevoflurane exposure. (E) Summary data showing decreased cerebral infarct size and improved neurological score in WT mice but not *Atf5*-cKO mice, n = 5-7 per group. (F) mRNA expression levels of the UPR^mt^-related genes in the cortices of WT mice after exposure to sevoflurane for 6 hours, n = 4-9 per group. (G) mRNA expression levels of the UPR^mt^-related genes in the cortices of *Atf5*-cKO mice after exposure to sevoflurane for 6 hours ,n = 5 per group. (H) Left: Mitochondrial function, measured as mitochondrial OCR, determined by assessing respiration of mitochondria isolated from the cortices of mice in control and sevoflurane groups (n = 5 per group). ADP, oligomycin, CCCP, and antimycin were added sequentially, as indicated by arrows. Right: Quantification of OCR after excluding non-mitochondrial respiration, n = 5 per group. Values are presented as means ± SD (n.s., not significant; *p < 0.05, **p < 0.01, ***p < 0.001).

### Sevoflurane induces mitochondrial dysfunction in *Atf5*-cKO mice

ATF5 promotes UPR^mt^ by increasing the expression of genes necessary for overcoming mitochondrial dysfunction and maintain homeostasis. To evaluate the potential role of ATF5-dependent UPR^mt^ in sevoflurane-induced preconditioning, we next evaluated sevoflurane-induced changes in the expression of UPR^mt^-related genes in *Atf5*-cKO mice 6 hours after sevoflurane exposure. In contrast to WT mice, in which various genes involved in UPR^mt^ were significantly upregulated after sevoflurane exposure (Fig. 2 F), *Atf5*- cKO mice showed no increase in UPR^mt^-related genes (Fig. 2 G). Importantly, mitochondrial respiration in isolated mitochondria was significantly decreased 24 hours after sevoflurane exposure only in *Atf5*-cKO mice (basal and state3u, Fig. 2 H). Our results imply that sevoflurane-induced ATF5 expression plays an important role in inducing UPR^mt^ and maintaining mitochondrial function after sevoflurane exposure in the brain.

### Sevoflurane-induced expression of GDF15 mediates preconditioning and mitochondrial upregulation

During stressful conditions, mitochondria release stress-responsive cytokines, also known as mitokines, which act as downstream modulators of UPR^mt^ (Jena *et al*, 2023; Kang *et al*, 2021). Growth differentiation factor 15 (GDF15), a representative mitokine that is widely expressed in the brain (Schindowski *et al*., 2011), has been shown to be involved in cardiac preconditioning (Kempf *et al*, 2006). Interestingly, we also found a significant increase in *Gdf15* mRNA levels in the cerebral cortex 6 hours after sevoflurane exposure in WT mice but not in *Atf5*-KO mice (Fig. 2 F, G). These results suggest that GDF15 may function downstream of ATF5 in inducing preconditioning in the brain. To further investigate the role of GDF15, we exposed *Gdf15*-KO mice to sevoflurane. Consistent with results obtained in *Atf5*-cKO mice, sevoflurane-induced preconditioning did not develop in *Gdf15*-KO mice, as reflected in the absence of a change in infarct size or improvement in neurologic scores (Fig. 3 A, B). Measurements of changes in the expression of mitochondrial energy metabolism-related genes showed that, whereas 18 of 83 genes related to mitochondrial energy metabolism were increased in WT mice, none of these genes were increased after sevoflurane exposure in the cerebral cortex of *Gdf15*-KO mice (Fig. 3 C, Suppl. Tables 1, 2). Furthermore, similar to the case in *Atf5*-cKO mice, sevoflurane exposure significantly decreased mitochondrial respiration in mitochondria isolated from the cerebral cortex of *Gdf15*-KO mice 24 hours after sevoflurane exposure (Fig. 3 D, E).

**Figure 3.**
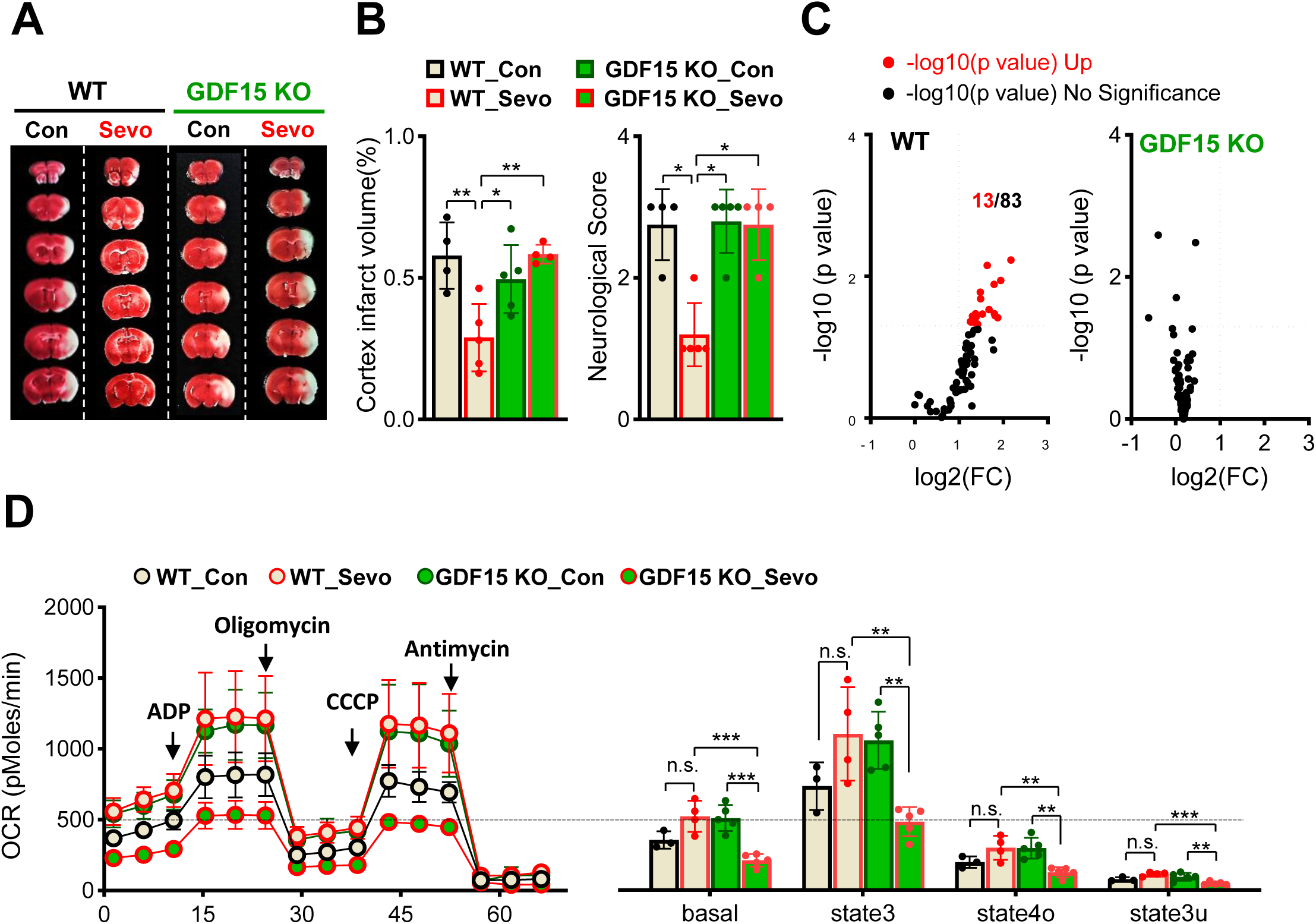
Sevoflurane exposure does not induce preconditioning in *Gdf15*-KO mice but does induce mitochondrial dysfunction. (A) Representative TTC-stained images of MCAO model WT and *Gdf15*-KO mice after sevoflurane (sevo) exposure. (B) Summary data showing decreased cerebral infarct volume, n = 4-5 per group. (C) Volcano plots showing mitochondrial energy metabolism-related genes whose levels were increased in the cerebral cortex of WT and *Gdf15*-KO mice 6 hours after exposure to sevoflurane, n = 3-5 per group. Mitochondrial energy metabolism-related genes are listed in Suppl. Table 1, 2. (D) Left: Mitochondrial function, measured as mitochondrial OCR, determined by assessing respiration of mitochondria isolated from the cortices of mice in control and sevoflurane groups. ADP, oligomycin, CCCP, and antimycin were added sequentially, as indicated by arrows. Right: Quantification of OCR after excluding non-mitochondrial respiration, n = 3-5 per group. Values are presented as means ± SD (n.s., not significant; *p < 0.05, **p < 0.01,***p < 0.001).

### ATF5 overexpression enhances GDF15 expression and mitochondrial function in excitatory neurons

Our results suggest that ATF5 regulates GDF15 expression. Although a direct relationship between ATF5 and GDF15 has not been previously established, previous studies have shown that GDF15 expression is downstream of ATF4, another major transcription factor of the UPR^mt^ (Jena *et al*., 2023; Kang *et al*., 2021). Considering that ATF5 expression is regulated by ATF4 (Juliana *et al*, 2017; Teske *et al*, 2013; Zhou *et al*, 2008), it is possible that GDF15 expression is also regulated through ATF5. To confirm that GDF15 is regulated by ATF5, we constructed a viral vector overexpressing ATF5 specifically in excitatory neurons (pAAV-hSyn-DIO-ATF5). AAVs containing the construct were injected into the cerebral cortex of Emx1**^cre/+^**mice (Fig. 4 A) and were found to increase ATF5 expression after 4 weeks (Fig. 4 B, Suppl. Fig. 3). Consistent with our initial results, ATF5 overexpression was associated with increased expression of genes (33 of 84) related to mitochondrial energy metabolism, including *Gdf15* (Fig. 4 C, Suppl. Table 3). Mitochondrial respiration was also increased in mitochondria isolated from cerebral cortices expressing AAVs (Fig. 4 D). Our results provide direct evidence that expression of GDF15, which plays a significant role in regulating mitochondrial function, is regulated by ATF5 in excitatory neurons.

**Figure 4.**
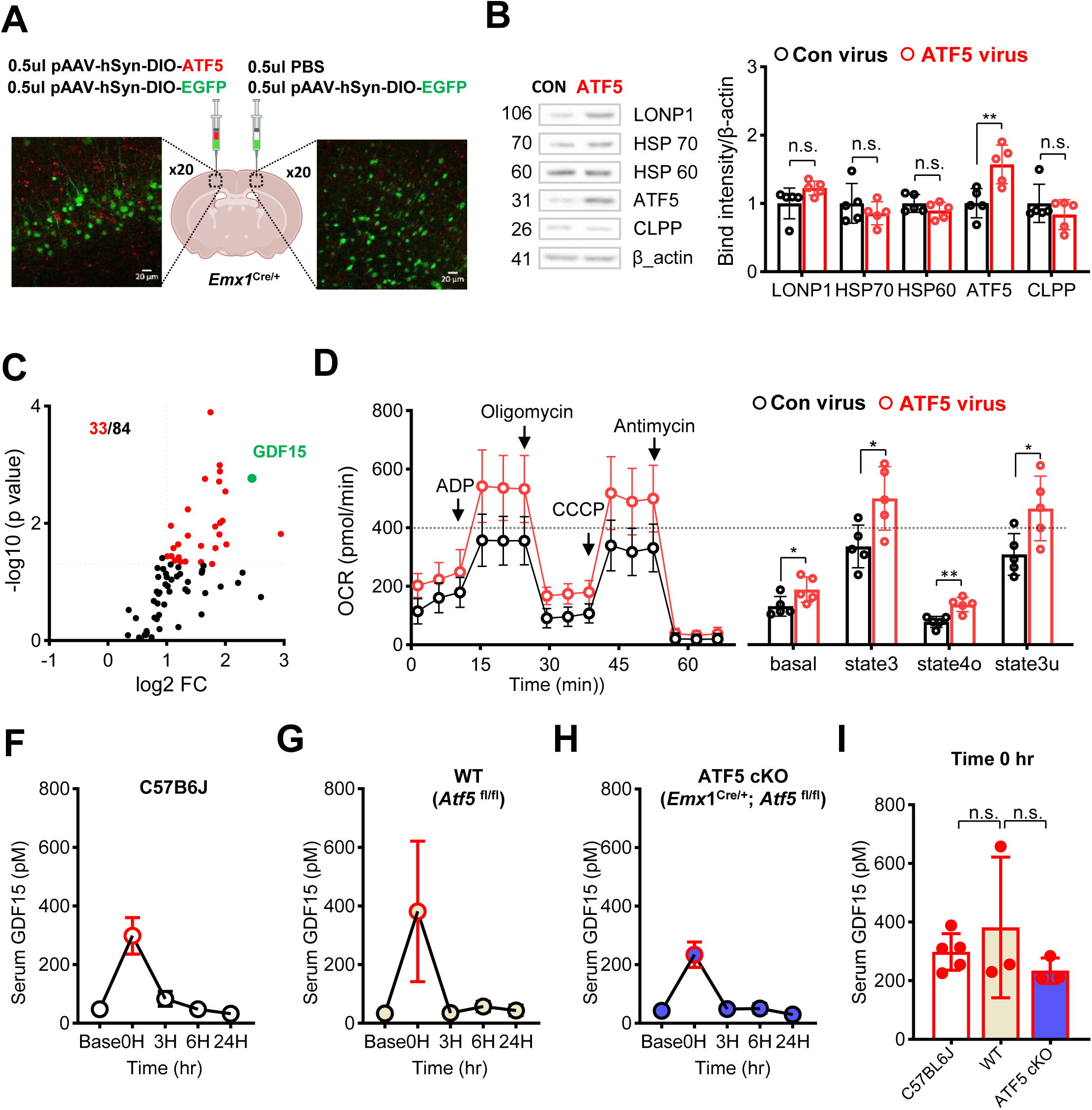
Overexpression of ATF5 increases GDF15 levels and mitochondrial function in the cortex, whereas serum GDF15 levels returned to baseline within 6 hours after exposure to sevoflurane. (A) Representative IHC images confirming pAAV-hSyn-DIO-ATF5 expression after stereotaxic injection of pAAV-hSyn-DIO-ATF5 (red) and pAAV-hSyn-DIO-EGFP (green) into the cortex of EMX1^Cre/+^ mice. (B) Western blot analysis of cortex surrounding the injection region. Cortex samples obtained 24 hours after sevoflurane exposure were analyzed for expression of the proteins, n = 5 per group. (C) Volcano plots showing mitochondrial energy metabolism-related genes (including GDF15) whose levels were increased around the injection region (n = 5 per group). Mitochondrial energy metabolism-related genes are listed in Suppl. Table 3. (D) Left: Mitochondrial function, measured as mitochondrial OCR, determined by assessing respiration of mitochondria isolated from the cortex of mice in control virus and ATF5 virus groups. ADP, oligomycin, CCCP, and antimycin were added sequentially, as indicated by arrows. Right: Quantification of OCR after excluding non-mitochondrial respiration, n = 5 per group. (F-H) Serum GDF15 levels prior to and 3, 6, and 24 hours after exposure to sevoflurane in C57BL/6J, WT and *Atf5*-cKO mice. (I) Comparison of serum GDF15 in the absence of sevoflurane exposure in C57BL/6J, WT and *Atf5*-cKO mice (n = 3-5). Full Western blot images are provided in Suppl. Fig 3. Values are presented as means ± SD (n.s., not significant; *p < 0.05, **p < 0.01,***p < 0.001).

Compared with the brain, other organs, such as the liver and kidney, express significantly higher levels of GDF15 (Yi, 2019). Therefore, it is possible that sevoflurane increases systemic GDF15 through these organs, which may act on the brain. Indeed, we observed an immediate increase in serum GDF15 levels following sevoflurane exposure in C57BL/6J, *Atf5*^fl/fl^ (WT), and EMX1^Cre/+^;*Atf5*^fl/fl^ (*Atf5*-cKO) mice (Fig. 4 F-H). Although serum levels returned to baseline within 6 hours, our results suggest that sevoflurane exposure alone is capable of increasing GDF15 in multiple organs, leading to an immediate rise in systemic GDF15. However, considering that sevoflurane-induced preconditioning did not develop in *Atf5*-cKO mice despite the significant increase in serum GDF15 levels, our results suggest that only ATF5-dependent GDF15 endogenously released in excitatory neurons is required for preconditioning. Furthermore, there was no difference in serum GDF15 levels among C57BL/6J, WT and *Atf5*-cKO mice (Fig. 4 I), suggesting that GDF15 expressed in excitatory neurons does not contribute to the increase in serum GDF15 after sevoflurane exposure.

### Sevoflurane-induced activation of ATF5-dependent UPR^mt^, upregulation of GDF15, and mitochondrial protection are absent in the cerebral cortex of aged mice

Unlike preclinical studies, which consistently report anesthesia-induced preconditioning, few clinical studies have suggested the possibility of neuroprotection after anesthesia (Lomivorotov *et al*., 2022; Raub *et al*., 2021). Since we found that ATF5-dependent UPR^mt^ activation, GDF15 expression, and mitochondrial upregulation are key players in sevoflurane-induced preconditioning, it is possible that these pathways are altered by specific factors in clinical trials. One potential factor is the age of patients. Whereas preclinical studies have been performed in young mice, most clinical studies have been performed in elderly patients (Chu *et al*, 2015; Guay *et al*, 2016; Memtsoudis *et al*, 2013; Neuman *et al*, 2021). Moreover, despite the fact that previous studies suggest that the inducibility of UPR may decline with aging (Munoz-Carvajal & Sanhueza, 2020), the effects of aging on sevoflurane-induced UPR^mt^ activation in the brain have not been studied in detail. To investigate the impact of aging on sevoflurane-induced preconditioning, we exposed aged mice (18-20 months) to sevoflurane. Compared to the case in young mice (Fig. 3 C), sevoflurane exposure increased very few genes related to mitochondrial energy metabolism in aged mice (Fig. 5 A, Suppl. Table 4). There was also no increase in the expression of UPR^mt^ proteins 24 hours after sevoflurane exposure (Fig. 5 B, C & Suppl. Fig. 4). Importantly, and similar to results obtained in *Atf5*-cKO and *Gdf15*- KO mice, mitochondrial respiration (OCR) was significantly decreased in isolated mitochondria from aged mice 24 hours after sevoflurane exposure (Fig. 5 D). Unfortunately, we were unable to determine whether sevoflurane-induced preconditioning was also absent in aged MCAO model mice because none of the aged mice survived 24 hours after performing the MCAO procedure.

**Figure 5.**
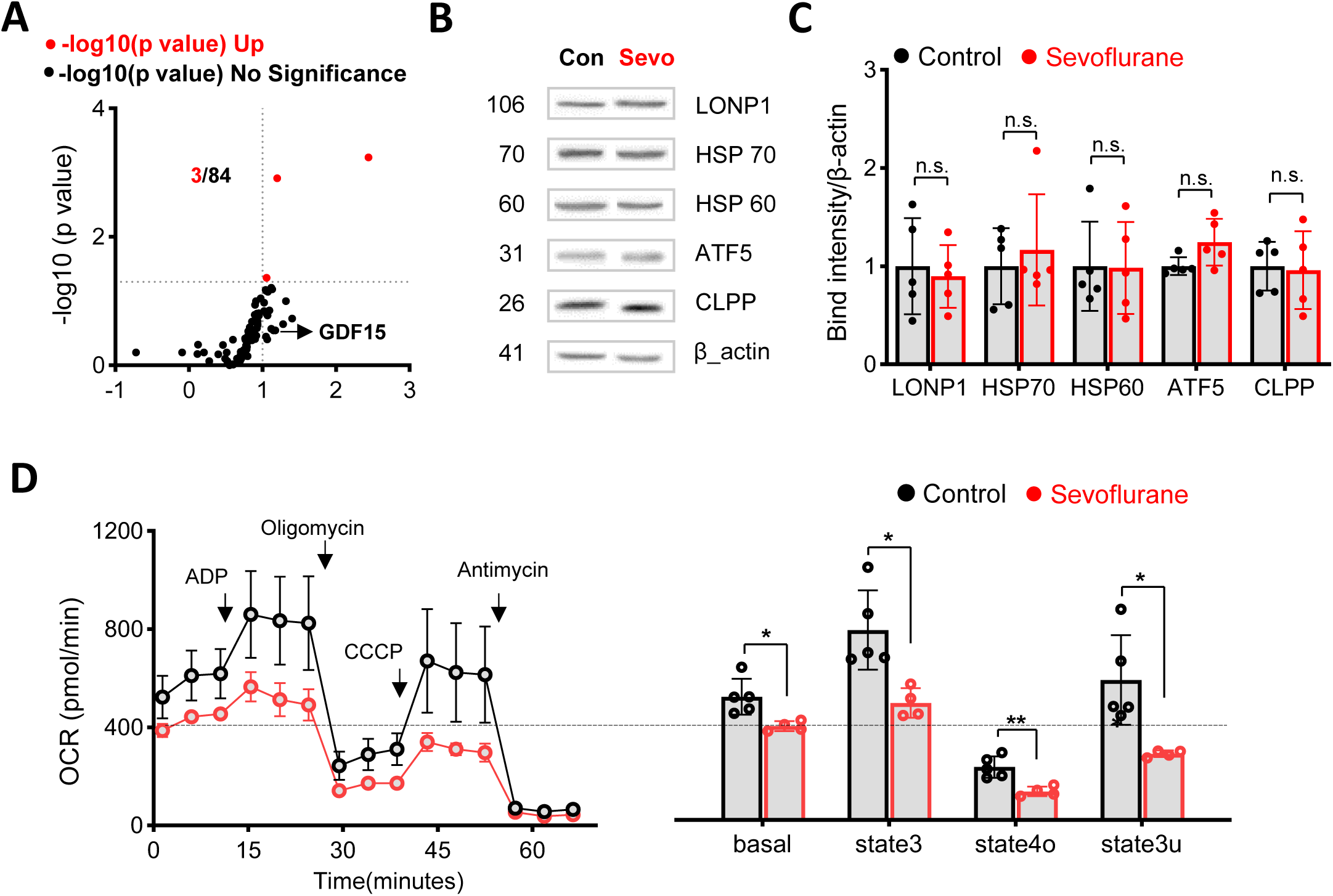
Sevoflurane-induced activation of ATF5-dependent UPR^mt^, GDF15 expression, and mitochondrial protection are absent in the cerebral cortices of aged mice. (A) Volcano plots showing mitochondrial energy metabolism-related genes whose levels were increased in the cerebral cortices of aging mice 6 hours after exposure to sevoflurane (n = 3-4 per group). Mitochondrial energy metabolism-related genes are listed in Suppl. Table 4. (B,C) Western blot analysis of proteins in cortical samples from aging mice obtained 24 hours after sevoflurane exposure, n = 5 per group. (D) Left: Mitochondrial function, measured as mitochondrial OCR, determined by assessing respiration of mitochondria isolated from the cortices of aging mice in control and sevoflurane groups, n = 4-5 per group. ADP, oligomycin, CCCP, and antimycin were added sequentially, as indicated by arrows. Right: Quantification of OCR after excluding non-mitochondrial respiration, n = 5 per group. Full Western blot images are provided in Suppl. Fig 4. Values are presented as means ± SD (n.s., not significant; *p < 0.05).

## Discussion

Neuroprotective strategies may be beneficial in aging societies to reduce the severity of perioperative stroke. In this study, we demonstrated that sevoflurane-induced neuroprotection is associated with the upregulation of genes involved in UPR^mt^ and mitochondrial metabolism. Our findings emphasize the critical role of ATF5, a key transcription factor, in mediating these protective effects. Specifically, we observed that sevoflurane preconditioning significantly upregulates ATF5 and its downstream target GDF15 – a regulator of mitochondrial function – in the cerebral cortex. This upregulation is critical for the neuroprotective effects following ischemic injury. To our knowledge, this is the first demonstration that sevoflurane-induced preconditioning is mediated by ATF5-regulated GDF15 expression in the brain. Notably, we found that this mechanistic pathway was not activated in the brain of aged mice, suggesting that age-specific strategies may be necessary for reducing the risk of perioperative stroke.

ATF5 plays a significant role in maintaining mitochondrial function, particularly in the context of cellular stress responses. We further found that the upregulation of ATF5 was necessary for the development of sevoflurane-induced preconditioning, as evidenced by the failure of sevoflurane exposure to improve infarct size or neurological outcomes in *Atf5*-cKO mice. The role of ATF5 in regulating mitochondrial function and inducing preconditioning after sevoflurane exposure is also evident through a comparison of changes in gene expression following sevoflurane exposure and viral overexpression of ATF5. Among the 83 mitochondrial energy metabolism-related genes examined, 19 were upregulated after sevoflurane exposure, 11 of which overlapped with genes increased following ATF5 overexpression in the cortex (Suppl. Table 5). Our findings are consistent with previous studies that identified ATF5 as a major regulator of UPR^mt^, where it governs the expression of genes essential for maintaining mitochondrial integrity under stress conditions (Wang *et al*, 2023b; Wang, 2019).

A significant finding of this study is the identification of GDF15 as a downstream effector of ATF5 in excitatory neurons. GDF15, a member of the transforming growth factor-beta (TGF-β) superfamily, is associated with various pathological conditions (Moon *et al*, 2020; Nga *et al*, 2021; Rochette *et al*, 2020; Siddiqui *et al*, 2022). Previous rodent studies have shown that treatment with GDF15 can reduce weight by increasing insulin sensitivity and enhancing energy expenditure through the GFRAL (glial cell line-derived neurotrophic factor [GDNF] family receptor alpha-like) receptor in the hindbrain (Sjøberg *et al*, 2023; Wang *et al*, 2021; Wang *et al*, 2023a). GDF15 is also upregulated in response to ischemic and hypertrophic conditions in the heart and liver, where it acts as a protective mechanism to prevent cellular damage (Àurea Navarro-Sabaté, 2006; Jian Xu, 2006; Kempf *et al*., 2006). While a previous study suggested that GDF15 provides protection against ischemia-reperfusion injury by inhibiting inflammation, particularly neutrophil infiltration and migration (Zhang *et al*, 2016), its role in the brain, especially in the context of anesthetic preconditioning, remains unclear. Our results also suggest GDF15 as a potential therapeutic agent that may provide protection against ischemia/reperfusion injury. However, unlike the case in previous studies (Chiariello *et al*, 2022), our data suggest that sevoflurane-induced preconditioning is mediated by endogenous GDF15 expressed in neurons, rather than by systemic GDF15 via the GFRAL receptor. In fact, preconditioning failed to develop in *Atf5*-cKO mice, despite a significant increase in serum GDF15 in these mice. Therefore, rather than directly targeting GDF15, therapies aimed at activating UPR^mt^ to maintain mitochondrial homeostasis may be more effective in the brain. It should be further noted that GDF15 has also been considered an inverse marker of general health, as its levels are significantly elevated in many chronic pathological conditions (Tsai *et al*, 2018). One recent study reported that treatment with ponsegromab, a humanized monoclonal antibody that inhibits GDF15, reduced cachexia symptoms in cancer patients with elevated serum GDF15 levels. Although our results suggest that serum GDF15 levels do not influence sevoflurane-induced preconditioning in the brain, further studies are needed to evaluate the effects of increased serum GDF15 levels after sevoflurane exposure (Dogon *et al*, 2024).

Another key finding of our study is the lack of sevoflurane-induced upregulation of ATF5 in aged mice. While the idea of providing protection against ischemia/reperfusion injury by simply applying anesthesia initially seemed uncontroversial, studies soon suggested that such protection may not occur in the aged heart (Boengler *et al*, 2009). Although most preclinical and clinical studies in older individuals have focused on cardiac protection, such studies may provide significant insights into brain preconditioning since the heart and brain share underlying preconditioning mechanisms (Chen *et al*, 2018; Xia *et al*, 2016). For instance, a previous study that compared young (3-5 months) and old (20-24 months) rats reported that isoflurane exposure was incapable of increasing reactive oxygen species levels or reducing myocardial size in aged cardiac tissue (Nguyen *et al*, 2008). A more recent study reported that the absence of sevoflurane preconditioning in aged rats (24 months) was attributable to the failure to activate NF-κB– regulated apoptotic genes (Zhong *et al*, 2017). In the current study, we provide another mechanism, showing that sevoflurane fails to activate ATF5-dependent UPR^mt^ in the brain of aged mice. Interestingly, high-intensity exercise (4 weeks) has been shown to activate UPR^mt^ in the skeletal muscles of 24-month-old mice (Cordeiro *et al*, 2021; Zhu *et al*, 2022). Although mitochondria from different organs may act differently, additional studies are necessary to show whether a prolonged or repeated stimulus can provide neuroprotection by activating UPR^mt^ in the aging brain.

There are several limitations of the present study. First, although we aimed to selectively knockout ATF5 in excitatory neurons (Emx1^cre/+^;*Atf5*^f/f^), ATF5 expression was only reduced by ∼30%. Thus, the presence of residual ATF5 expression complicates a full evaluation of ATF5’s complete function. Second, although we showed that sevoflurane exposure failed to activate ATF5-dependent UPR^mt^ in aged mice, we were unable to confirm the absence of sevoflurane-induced preconditioning, as none of the mice survived 24 hours after MCAO. However, given the similarity of preconditioning mechanisms in the brain and heart, and reports of absent preconditioning in the hearts of aged mice, it is likely that preconditioning also fails to develop in the brain. Third, the neuroprotective effect of sevoflurane was examined at a single time point (24 hours after sevoflurane exposure). Studies have shown that preconditioning develops in 2 phases: an early preconditioning phase that lasts for 2-3 hours immediately after anesthesia, and a late preconditioning phase that appears after 12-24 hours and lasts up to 72 hours (Lutz & Liu, 2006; Tsutsumi *et al*, 2006). Although it is possible that ATF5 and GDF15 also play significant roles in the early phase (Athiraman & Giri, 2024), additional studies are needed to confirm the mechanisms in each phase. Lastly, we have not evaluated the possible effects of increased serum GDF15 that occurred immediately after sevoflurane exposure. Previous studies have reported multiple roles of GDF15 through its actions on GFRAL, the currently known receptor for GDF15, in the brain (Breit *et al*, 2021).

In conclusion, we identified a novel mechanism of sevoflurane-induced preconditioning by manipulating the expression of the transcription factor ATF5 specifically in excitatory neurons. We demonstrated that sevoflurane exposure activates ATF5-dependent UPR^mt^, resulting in increased expression of genes related to mitochondrial function and energy metabolism, including GDF15, to maintain mitochondrial homeostasis.

Unlike most studies, which focus on systemic GDF15, our results suggest that endogenous GDF15 is essential for the development of neuroprotection against ischemia/reperfusion injury. Importantly, we also discovered that sevoflurane failed to activate the ATF5-GDF15 signaling pathway in the aged brain, leading to a decrease in mitochondrial function after sevoflurane. Considering that the age of patient populations is steadily increasing, therapeutic approaches that enhance mitochondrial function in the aged brain may provide additional protection against perioperative stroke.

## Materials and Methods

### Animals

This study was approved by the Committee on Animal Research at Chungnam National University (Daejeon, South Korea; 202203A-CNU-050). To control for fluctuations in estrogen levels, which can influence infarct size and neurological outcomes in female mice, only male mice were used. Mice were housed in cages (<6 mice/cage) at 24-25°C under a 12-hour light/dark regimen and provided food and water *ad libitum*. For creation of conditional KO mice, floxed *Atf5* (*Atf5*^fl/fl^) mice in which exon 3 was targeted were first generated (Cyagen Biosciences, CA, US) and backcrossed with C57BL/6J mice (Damul Science, Daejeon, Korea) for more than three generations. *Atf5*^fl/fl^ mice were then crossed with Emx1^cre^ knock-in mice to produce Emx1^cre^;*Atf5*^fl/fl^ conditional KO (*Atf5*-cKO) mice (Jessica A. Gorski, 2002; Takuji Iwasato, 2004). GDF15-KO mice were derived from an inbred C57BL/6 strain that was kindly provided by Dr. S. Lee (Johns Hopkins University School of Medicine, Baltimore, MD, USA).

### Sevoflurane exposure

Mice were divided into two groups: Control and Sevoflurane. Mice in the Control group were placed in an anesthesia chamber and exposed to a constant flow of fresh gas (fraction of inspired oxygen [FiO_2_], 0.4 at 4 L/min) for 2 hours and 10 minutes. Mice in the Sevoflurane group were treated identically, but received 2.5% sevoflurane added to the fresh gas for 2 hours, followed by 10 minutes without sevoflurane for recovery. The body temperature of mice was maintained by placing the anesthesia chamber in a 37°C water bath. FiO_2_ and sevoflurane concentrations were monitored using an m-CAiO gas analyzer module (Datex-Ohmeda, Helsinki, Finland). Adequate respiration was confirmed by blood gas analysis in a subset of mice (Table 1).

### Blood gas analysis

To confirm that our protocol did not cause significant respiratory distress, we performed a blood gas analysis on arteriovenous mixed blood collected by decapitation directly after sevoflurane exposure (Chung *et al*, 2017). Control group mice also were briefly exposed to sevoflurane for ethical reasons. Mixed blood was analyzed using iSTAT (Abbott Park, Illinosis, USA).

### MCAO model

The MCAO model was generated as previously described (Chiang *et al*, 2011). Tracheal intubation was performed after sevoflurane exposure (5%, 3 minutes) using a 20-gauge intravenous catheter (REF 382434; BD Angiocath Plus, Singapore). To ensure proper positioning of the catheter so as to avoid one-lung ventilation and gas leakage during ventilation, we modified the intravenous catheter using the distal portion of a 200 μl pipette tip (Suppl. Fig. 5 A). Mice were mechanically ventilated (Minivent Model 845; Havard Apparatus) with 2.5% sevoflurane (FiO_2_, 40% at 4 L/min; respiratory rate, 180/min; tidal volume, 10 μl/g). Rectal temperature was kept at 37°C ± 0.5°C using a heating pad. A blood gas analysis was performed to confirm adequate respiration (Suppl. Fig. 5 B). After exposing the right common carotid artery (CCA) via a ventral midline neck incision, the right external carotid artery (ECA) and internal carotid artery (ICA) were identified. The CCA and ECA were permanently clamped, and a 6-0 silicone rubber-coated monofilament (L56PK10; Doccol Corporation, Sharon, MA, USA) was carefully inserted into the right CCA, upstream from the ECA clamp, and advanced towards the ICA until slight resistance was felt. The monofilament was inserted into the artery from the bifurcation of the ECA and ICA to the MCA, a distance of approximately 9-10 mm. The suture was then tightly fastened around the CCA for 40 minutes. After the occlusion period, the suture was removed, and the CCA was permanently fastened to prevent blood flow. Changes in microvascular blood flow in the MCA territory (1 mm posterior and 5 mm lateral to Bregma on the right parietal cranial region) were confirmed by laser-Doppler microscopy (PeriFlux 6000; PERIMED, Sweden). Only mice with ≥ 80% flow reduction during the ischemic period were included in this study (Saeed Ansari1, 2011).

### Evaluation of neurological function and infarct volume in the MCAO model

The behavior of MCAO mice was evaluated using a modified Bederson scoring method, as follows: 0, no deficit; 1, forelimb flexion; 2, forelimb flexion plus decreased resistance to lateral push; 3, unidirectional circling; 4, longitudinal spinning or seizure activity; 5, no movement (Bederson *et al*, 1986). Infarct size was measured using 2,3,5-triphenyltetrazolium chloride (TTC) staining. Briefly, 24 hours after MCAO surgery, coronal brain slices (1000 mm) were obtained using a VT1200 vibratome (Leica Microsystems, Wetzlar, Germany). The brain slices were placed in 2% TTC (Sigma-Aldrich, St. Louis, MO, USA) in phosphate-buffered saline (PBS) for 20-25 minutes at room temperature (RT) and then fixed in 10% neutral buffered formalin solution (Sigma-Aldrich) at 4°C until imaging. The infarct size was calculated using the ImageJ program.

### Western blotting

Protein samples from cerebral cortex tissue were obtained 24 hours after sevoflurane exposure by homogenizing the cortex in lysis buffer containing phosphatase inhibitors (Sigma-Aldrich, Cat# PHOSS-RO) and protease inhibitors (Sigma-Aldrich, Cat# P8340) using TissueLyser II (Qiagen, Germany). The homogenate was then centrifuged at 12,000 × g for 20 minutes at 4°C and supernatants were collected. Samples from primary cultured neurons (see below) were obtained by first aspirating the culture media and washing the cells with ice-cold PBS, then lysing cells in a buffer containing phosphatase inhibitors and protease inhibitors. Protein concentration in tissue homogenates and cultured cell lysates was measured using the SMART BCA Protein Assay Kit (iNtRON). Proteins (20 μg/sample) were resolved by sodium dodecyl sulfate-polyacrylamide gel electrophoresis (SDS-PAGE; Bio-Rad, CA, USA) and transferred to polyvinylidene fluoride (PVDF) membranes (0.45 μm pore size; Merck Millipore Ltd., Ireland) at 385 mA for 1 hour. Membranes were blocked with 5% skim milk in Tris-buffered saline containing Tween 20 (TBS-T; 10 mM Tris-HCl, pH 7.6, 150 mM NaCl, 0.1% Tween 20) for 1 hour, followed by incubation with primary and appropriate secondary antibodies. The indicated primary antibodies against the following proteins were used: CLPP (Abcam, CAT# ab124822), ATF5 (Abcam, CAT# ab184923), phosphorylated-Eif2α (p-Eif2α; Cell Signaling Technology, CAT#3398), Eif2α (Cell Signaling Technology, CAT#5324ATF4), ATF4 (Abcam, CAT# ab184909), HSP60 (Abcam, CAT# ab46798), HSP70 (Santa Cruz Biotechnology, CAT#J1216), ATF6 (Abcam, CAT# ab203119), BIP (GRP78) (Abcam, CAT#ab108615), LONP1 (Abcam, CAT#ab103809), p-IRE1 (S724) (Abcam, CAT# ab124945), IRE1 (Abcam, CAT# ab37037), total PERK (t-PERK; Cell Signaling Technology, CAT#C33E10), p-PERK (Thr982, Affinity Biosciences, CAT#DF7576), and β-actin (Santa Cruz Biotechnology, CAT# sc-8432).

### Oxygen consumption rate

Mitochondria were isolated from the cerebral cortex 24 hours after exposure to sevoflurane as previously described (Lee *et al*, 2024). In brief, the cerebral cortex was homogenized in mitochondrial isolation buffer (70 mM sucrose, 210 mM mannitol, 5 mM HEPES, 1 mM EGTA, and 0.5% [w/v] fatty acid-free BSA [pH 7.2]) using a Teflon-glass homogenizer (Thomas Scientific, USA). Following centrifugation at 600 × g for 10 minutes at 4°C, and at 8,000 × g for 10 minutes at 4°C, the mitochondrial fraction was resuspended in mitochondrial isolation buffer. Protein concentration was determined using the Bradford assay (Bio-Rad). Aliquots containing 2ug/25ul or 20 µg/50ul of protein were diluted with mitochondrial assay solution (70 mM sucrose, 220 mM mannitol, 10 mM KH_2_PO_4_, 5 mM MgCl_2_, 2 mM HEPES, 1 mM EGTA, 0.2% [w/v] fatty acid-free BSA, 10 mM succinate, and 2 μM rotenone [pH 7.2]) and seeded in Seahorse XFe96/XF Pro Cell Culture Microplates or XF-24 plates (Seahorse Bioscience, USA). The plates were centrifuged at 2,000 × g for 20 minutes at 4°C using a swinging-bucket microplate adaptor (Eppendorf, Germany). After adding 180ul or 450 µl of mitochondrial assay buffer, the plates were equilibrated at 37°C for 8-10 minutes and then transferred to the Seahorse XF pro or Seahorse XF-24 extracellular flux analyzer (Seahorse Bioscience). Mitochondrial respiration was assessed by measuring the oxygen consumption rate (OCR). Oxygen consumption measurements were obtained under five sequentially induced respiratory states: i) basal respiration; ii) State3 (coupled respiration), with addition of ADP (4 mM) for measuring oxygen consumption during ATP production; iii) state4o, with addition of oligomycin (2.5 μg/ml) for measuring oxygen consumption from protein leakage; iv) state3u (uncoupled respiration), with addition of carbonyl cyanide m-chlorophenyl hydrazine (CCCP, 4 μM) for measuring oxygen consumption during maximal respiration; and v) non-mitochondrial respiration, with addition of antimycin (4 μM). OCR was automatically calculated and recorded using the Seahorse wave pro or Seahorse XF-24 software (Seahorse Bioscience).

### Real-time polymerase chain reaction

Cortex samples were collected 6 hours after sevoflurane exposure. Total RNA was extracted, and cDNA was synthesized using a High-Capacity cDNA Reverse Transcription Kit (Invitrogen, CA, USA). Cortical mRNA levels were quantified by quantitative reverse-transcription polymerase chain reaction (RT-qPCR), performed on a real-time Exicycle system (BIONEER, Daejeon, Korea) using 100 ng of cDNA, 2X SYBR mix, and forward and reverse primers (3 pmol each). An AccuTarget qPCR Screening Kit (BIONEER, Daejeon, Korea) was used to assess the expression of 83 genes related to mitochondrial energy metabolism. Expression of the following genes involved in UPR^mt^ were measured using the indicated primer pairs: *Lonp1*, 5’-GAC CAT TCC GGG ATA TCA TCG-3’ (forward) and 5’-CGATGATATCCCGAATGGTC- 3’ (reverse); *mtDNAj*, 5’-GGA TAG GCG AGA GGC TGG-3’ (forward) and 5’-CCA GCC TCT CGC CTA TCC-3’ (reverse); *Atf5*, 5’-CAA GGA TCC TCG GAT CTT-3’ (forward) and 5’-AAG GCG AAG GTG GAG GAC-3’ (reverse); *Hsp60*, 5’-AAA TGC TTC GAC TAC CCA CAG-3’ (forward) and 5’-CTG TGG GTA GTC GAA GCA TTT-3’ (reverse); *Hsp10*, 5’-AAG TTT CTT CCG CTC TTT GACA-3’ (forward) and 5’-TGT CAA AGA GCG GAA GAA ACT T-3’ (reverse); *Clpp*, 5’-GAG TCA GCA ATG GAG AGG GA-3’ (forward) and 5’- TCC CTC TCC ATT GCT GAC TC-3’ (reverse); and *GDF15*, 5’-AGG ACC TGC TAA CCA GGC TG-3’ (forward) and 5’-TCT GGC GTG AGT ATC CGG AC-3’ (reverse). Because there are two splice variants of *Atf5*, we separately measured mRNA levels of total *Atf5*, *Atf5*-α, and *Atf5*-β to confirm the lack of *Atf5* gene expression in *Atf5*-cKO mice (Torres-Peraza *et al*, 2013): total *Atf5*, 5’-GTC TTC ACC CAG CTG AAC AAT- 3’ (forward) and 5’-AAT GGA GGC TGC ACC AAC-3’ (reverse); *Atf5ɑ*, 5’-GTT GCC TCC TCG CCT TTT-3’ (forward) and 5’-GGA GGC TGC ACC AAC AAT-3’ (reverse); and *Atf5β*, 5’-TTT TAT GAA GAG GAA TAA GAT GAG GTC-3’ (forward) and 5’-GGA GGC TGCA CCA ACA AT-3’ (reverse). mRNA expression levels were normalized using β-actin as a housekeeping gene: *β-actin*, 5’-GGC TGT ATT CCC CTC CAT CG-3’ (forward), and 5’-CCA GTT GGT AAC AAT GCC ATG T-3’ (reverse).

### Immunohistochemistry (IHC)

Mice were cardiac perfused with PBS, followed by a 4% paraformaldehyde solution. Whole brains were extracted and preserved in 4% paraformaldehyde overnight at 4°C and then transferred to a 30% sucrose solution before further processing. Coronal sections (30-µm thick) were prepared and washed with PBS three times for 5 minutes each. Brain slices were blocked in a solution containing 3% Tween 20 and 2% bovine serum albumin (BSA) in PBS for 1 hour. Slices were rinsed with PBS and incubated overnight at 4°C with antibodies against CAMK II alpha (1:500; Santa Cruz) and ATF5 (1:500; Abcam), both diluted in the blocking solution. After washing with PBS three times for 5 minutes each, the sections were incubated with a fluorescently labeled secondary antibody for 2 hours at RT in the dark. Subsequently, sections were washed with PBS three times for 5 minutes each and incubated with DAPI (4′,6-diamidino-2-phenylindole, 1:1000 in PBS) for 15 minutes. Finally, the sections were washed with PBS three times for 5 minutes each and mounted with a suitable mounting solution. Slices were visualized using an LSM 980 confocal microscope (ZEISS, Oberkochen, Germany).

### Primary neuron culture

Cortical neurons were harvested from postnatal day 1 *Atf5*^fl/fl^ and Emx1^cre^;*Atf5*^fl/fl^ mice. The cortex was dissociated with papain, after which the resulting cell suspension was plated onto poly-D-lysine–coated, 18-mm glass coverslips in 60-mm Petri dishes containing plating medium, consisting of Neurobasal-A medium supplemented with 2% B-27, 10% FBS, 1% GlutaMax, and 1 mm sodium pyruvate (all from Thermo Fisher Scientific). After 4 hours, the plating medium was replaced with the same medium lacking FBS; thereafter, 50% of the medium was replaced every 7 days.

### Immunocytochemistry

Cells were fixed by aspirating the culture media, washing with ice-cold PBS, and adding a fixation buffer containing 4% formaldehyde and 4% sucrose in 1x PBS. After incubating at RT for 15 minutes, the cells were washed three times with ice-cold 1x PBS for 5 minutes each. Cells were permeabilized by incubating with 0.2% Triton X-100 in PBS at RT for 10 minutes. Cells were then washed three times with ice-cold 1x PBS and blocked for 1 hour at RT with a blocking solution containing donkey serum (GeneTex, Cat No.GTX30972). Primary antibody hybridization was carried out by incubating the primary antibody (diluted in blocking solution) overnight at 4°C. The following day, the primary antibody solution was removed, and cells were washed three times with ice-cold 1x PBS for 10 minutes each. For secondary antibody hybridization, cells were incubated with secondary antibody (diluted in blocking solution containing donkey serum) at 37°C for 2 hours in the dark. After washing three times with ice-cold 1x PBS (10 minutes each), cells were coverslip-mounted with Faramount mounting medium followed by sealing with clear nail polish. Slides were dried for 24 hours at RT in the dark prior to fluorescence microscopy analysis and stored at 4°C in dark chambers for long-term preservation.

### Virus packaging and stereotaxic brain injection

Full-length mouse *Atf5* (Cat #MR203843, Origene) was subcloned into pAAV-hSyn-DIO-EGFP (Addgene, plasmid #50457) to produce the pAAV-hSyn-DIO-ATF5 construct. AAVs (serotype PHP.eB) were packaged as previously described (Noh *et al*, 2022). The final virus solution was aliquoted and stored at -80°C. For stereotaxic brain injections, mice were anesthetized with 2.5% sevoflurane and then secured in a stereotaxic apparatus (RWD, Shenzhen, China). During the procedure, a continuous flow of 2.5% sevoflurane was maintained with fresh gas supplied at 4 L/min (FiO_2_, 0.4). Coordinates relative to Bregma were AP, -0.9 mm; ML, ±3.0 mm; and DV, -1.5 mm (Iraklis Petrof, 2014). AAVs were infused at a rate of 0.1 μl/min, with a post-injection diffusion time of 8-10 minutes. Mice were placed in a recovery chamber for at least 30 minutes after injection.

### Statistical analysis

R statistical software (version 4.2.0; R Core Team, Austria) was used for data analysis. All continuous variables were tested for normality and variance homogeneity. Independent t-test or one-way analysis of variance (ANOVA) was applied only if these two conditions were met. In cases where normality was not confirmed, the Kruskal–Wallis test was used. The Welch ANOVA was applied when variation lacked uniformity. P-values less than 0.05 were considered significant. Complete statistical results are provided in Supplementary Data. All data are presented as means ± standard deviation (SD).

## Acknowledgements

This work was supported by the National Research Foundation of Korea (NRF) grant funded by the Korea government (MSIT): NRF-2022R1A2C2006269 (RS-2022-NR070193); NRF-2022R1A2C2002756 (RS-2022-NR070073); RS-2024-00406568; RS-2024-00397681 and the Korea ministry of Health and Welfare: HR22C1734 (RS-2022-KH130308).

## Author contributions

Xianshu Ju, Tao Zhang: conceptualization; data curation; formal analysis; investigation; methodology; project administration; validation; visualization; writing-original draft. Jianchen Cui, Yulim Lee, Suho Lee, Ho Min Kim, Chul Hee Choi: investigation; methodology. Boohwi Hong: data curation, formal analysis, methodology. Hyon-Seung Yi, Jun Young Heo and Woosuk Chung: conceptualization; resources; supervision; funding acquisition; investigation; project administration; writing-review and editing.

