## Supplementary material for "ATF5-Dependent GDF15 Expression Mediates Anesthesia-Induced Neuroprotection Against Stroke": Suppl Figure

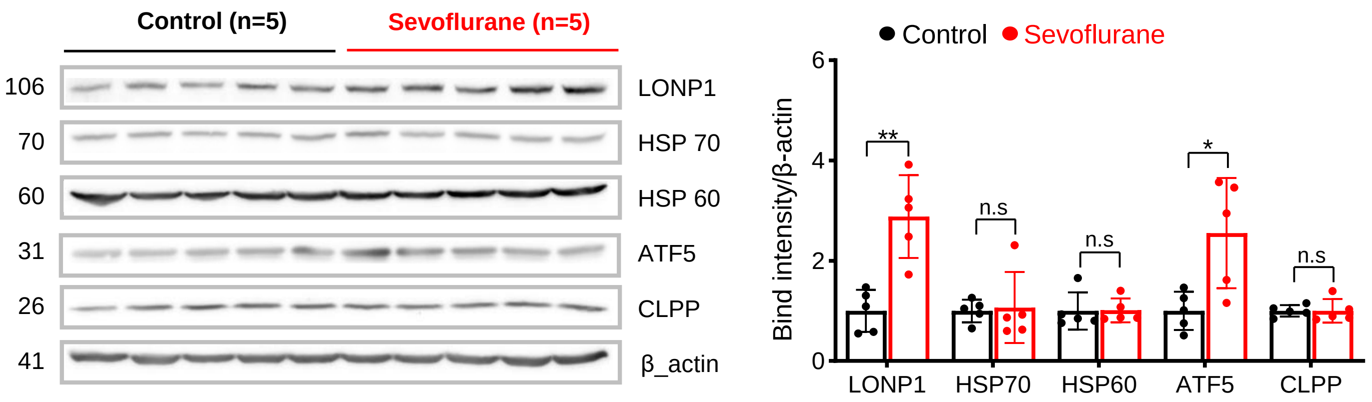


**Supplementary Figure 1. Full western blot images of figure 1E.**

Western Blotting of cortex samples obtained 24hr after sevoflurane exposure in 8weeks mice (n=5 per group).

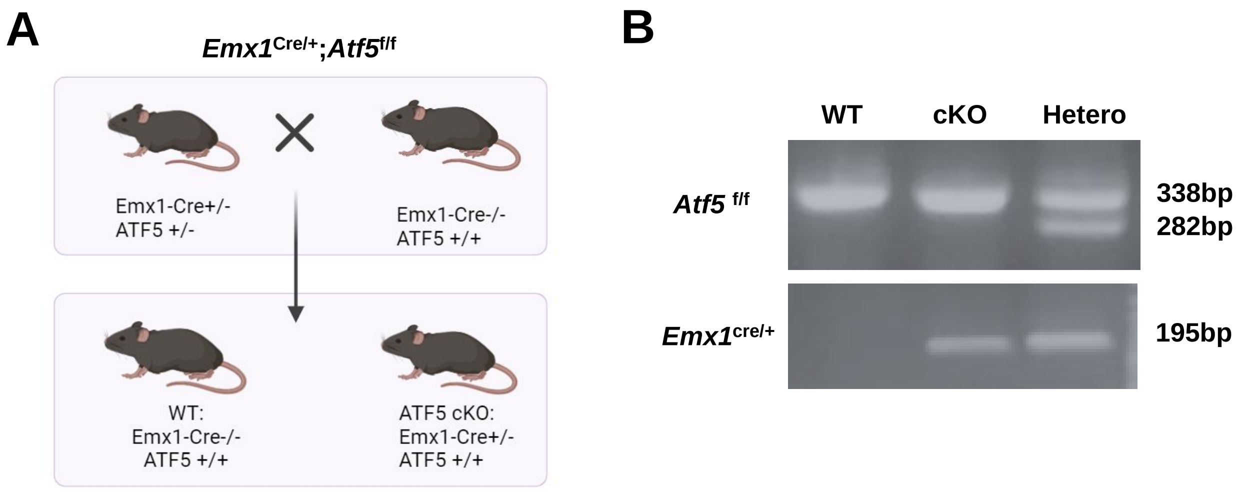


**Supplementary Figure 2. Generation of ATF5 conditional knockout mice.**

**(A)** Schematic drawing of the generation of ATF5 conditional knockout mice. **(B)** Genotyping results with two bands indicating heterozygous mice.

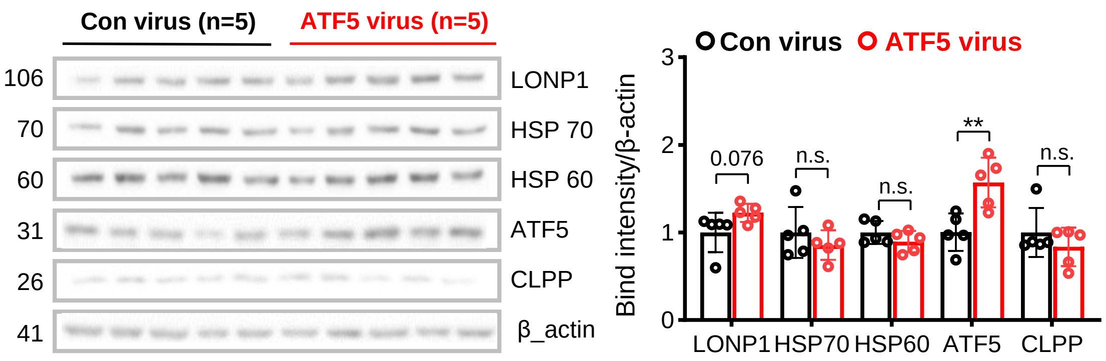


**Supplementary Figure 3. Full western blot images of figure 4B.**

Western Blotting of cortex samples obtained 4 weeks after virus injection (n = 5 per group).

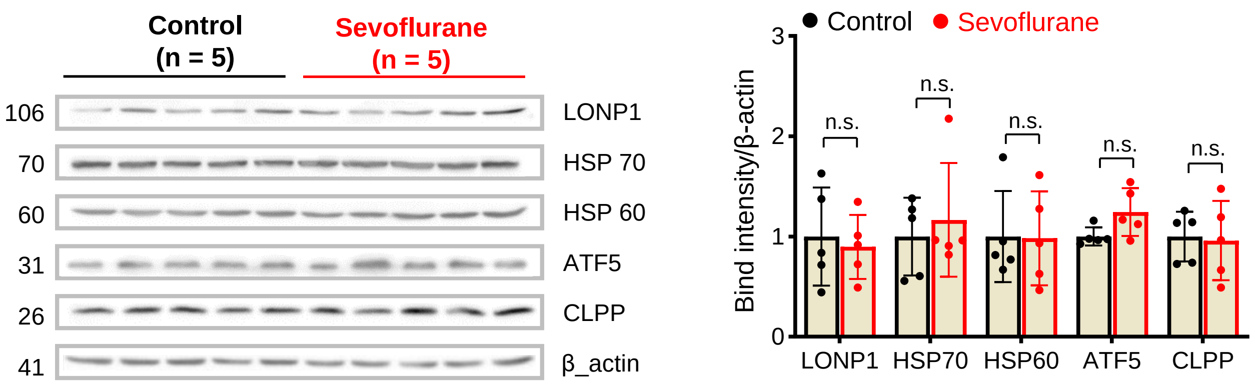


**Supplementary Figure 4. Full western blot images of figure 5B, C.**

Western Blotting of cortex samples obtained 24hr after sevoflurane exposure in aged mice (n=5 per group).

A

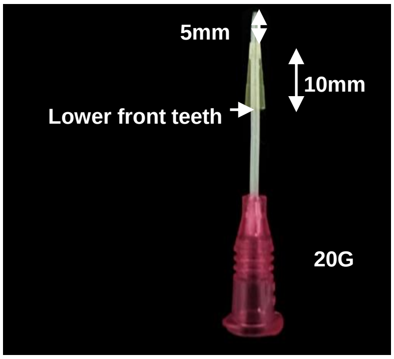


B

| | pH | $pCO_2$ mmHg | $pO_2$ mmHg | $sO_2$ % |
|---|---|---|---|---|
| **Animal 1** | 7.25 | 46.6 | 73 | 92 |
| **Animal 2** | 7.35 | 42.4 | 94 | 97 |
| **Animal 3** | 7.21 | 46.8 | 73 | 91 |
| **Animal 4** | 7.28 | 48.4 | 68 | 91 |
| **Mean±SD** | 7.27±0.05 | 46.05±0.22 | 77±10.02 | 92.75±2.49 |

**Supplementary figure 5. (A)** Customized endotracheal tube for mice. **(B)** Blood gas analysis using trunk blood obtained after decapitation with ventilation conditions used for MCAO surgery. pCO2, partial pressure of carbon dioxide; pO2, partial pressure of oxygen; sO2, oxygen saturation.
